# Integrated evolutionary and structural analysis reveals xenobiotics and pathogens as the major drivers of mammalian adaptation

**DOI:** 10.1101/762690

**Authors:** Greg Slodkowicz, Nick Goldman

## Abstract

Understanding the molecular basis of adaptation to the environment is a central question in evolutionary biology, yet linking detected signatures of positive selection to molecular mechanisms remains challenging. Here we demonstrate that combining sequence-based phylogenetic methods with structural information assists in making such mechanistic interpretations on a genomic scale. Our integrative analysis shows that positively selected sites tend to co-localise on protein structures and that positively selected clusters are found in functionally important regions of proteins, indicating that positive selection can contravene the well-known principle of evolutionary conservation of functionally important regions. This unexpected finding, along with our discovery that positive selection acts on structural clusters, opens new strategies for the development of better models of protein evolution. Remarkably, proteins where we detect the strongest evidence of clustering belong to just two functional groups: components of immune response and metabolic enzymes. This gives a coherent picture of immune response and xenobiotic metabolism as the drivers of adaptive evolution of mammals.

## Introduction

Over the course of evolution, the genomes of all organisms are shaped by the environment. The results of this process can be observed by comparing evolutionarily related sequences from different species: regions that code for essential cellular functions can remain unaltered over hundreds of millions of years, while changing evolutionary pressures can lead to emergence of new functions over very short evolutionary timescales. As a result, evolutionary histories of sites in the genome hold information about their functional importance. Functionally important regions are routinely identified by taking advantage of the fact that they are highly conserved in evolution ^1,2^. Similarly, methods for detecting regions harbouring adaptive changes have been developed to take advantage of the fact that rapid fixation of new alleles is a hallmark of positive selection ^3,4^. Analyses of patterns of evolutionary change can identify specific cases of adaptation as well as reveal general principles that guide evolution ^5^. Understanding evolutionary processes and distinguishing between neutral and adaptive changes is therefore one of the key aims of modern evolutionary studies.

As most proteins have to maintain a specific three-dimensional shape to perform their function, protein-coding genes exhibit particularly complex patterns of substitution. Biophysical constraints restrict the allowed amino-acid substitutions and result in dependencies across the entire protein sequence. While structural features can explain a significant proportion of observed site-to-site rate variation ^6^, previous studies have focused on evolutionary scenarios where existing functions are maintained and little is known about the structural properties of sites evolving under positive selection.

Present lack of understanding of structural aspects of adaptive evolution is particularly surprising bearing in mind that many single-gene studies took advantage of protein structure to assess the functional significance of positively selected sites identified from sequence data. In the classic study of Hughes and Nei ^7^, positively selected residues in the MHC molecule were found to cluster in the groove where pathogen-derived peptides are bound, supporting the hypothesis that rapid amino-acid substitutions at these sites tuned the ability to bind peptides derived from pathogens. Similarly, positively selected sites in TRIM5α, a viral restriction factor that can inhibit the cellular entry of HIV in non-human primates, are placed in the region that mediates binding to the virus ^8^. In these studies, as in others (e.g. ^9^), proximity of positively selected residues on the protein structure was used as corroborating evidence and helped assign a molecular mechanism underlying detected adaptations.

As the amount of available genomic data increased, studies of positive selection in individual proteins were followed by genome-wide positive selection scans ^10–15^. Such genomic scans, using appropriately adapted statistical methodology ^16,17^, can identify which cellular processes are primary targets of positive selection and generate testable hypotheses. However, structural aspects of identified examples were largely neglected and so no coherent view of how protein structure affects adaptive evolution has emerged from these investigations.

This is a significant gap in our understanding of evolution. Biophysical constraints restrict what substitutions are allowed for protein function to be maintained and are also likely to limit the emergence of adaptive changes in response to pressures from the environment, yet no evolutionary theory predicts the structural properties of sites harbouring adaptive changes. It is not established whether positive selection is more likely to act on protein sites where the effect of mutations is the largest (e.g. enzyme catalytic sites or key interaction interfaces) or regions where mutations likely have a smaller effect (e.g. allosteric regulation sites). Adaptive changes are associated with rapid fixation of advantageous mutations, yet functional regions are thought to be highly conserved in evolution. Contrasting these two principles leads to an apparent paradox.

Here, we integrated structural information into evolutionary analyses in order to study the properties of positively selected sites. We demonstrate that detailed mechanistic interpretation of findings can be achieved on a genome-wide level, just as in the case of earlier studies of individual proteins. In recent years, it has become apparent that structural data can be an orthogonal source of information that can serve to validate and augment findings in different areas of genomics ^18^. Structural placement of sites of interest, such as those identified through genome-wide sequence analyses, can be used to strengthen the confidence in findings — clustering of sites indicates concerted function whereas unrelated sites are expected to be more uniformly distributed in the structure. Recently developed methods based on clustering of sites on protein structures have been successful in distinguishing causal and hitchhiking mutations underlying genetic diseases ^19^ and for identifying mutations with functional impact in cancer ^20–23^. Detailed information about the protein structure can similarly aid understanding of molecular mechanisms underlying adaptation at detected sites.

To obtain a structurally-informed view of positive selection at the residue level, we developed an approach combining a genome-wide scan for positive selection with structural information (fig. 1a). We applied 3D clustering to detect genes with positively selected sites in a robust manner that additionally allowed us to link identified cases to an underlying molecular mechanism. We demonstrate that positively selected sites tend to occur close to one another on protein structure and detect 20 high-confidence positively selected clusters (table 1). Strikingly, we find that all but one of the identified cases are immune-related proteins or metabolic enzymes. In both of these functional categories, interactions with dynamic environmental parameters appear to have shaped the evolutionary histories of the genes involved. By further analysing the placement of positively selected clusters, we find that pervasive positive selection acts on regions that are typically highly conserved in evolution, suggesting new strategies for the development of more accurate models of protein evolution and methods for detecting positive selection.

**Figure 1.**
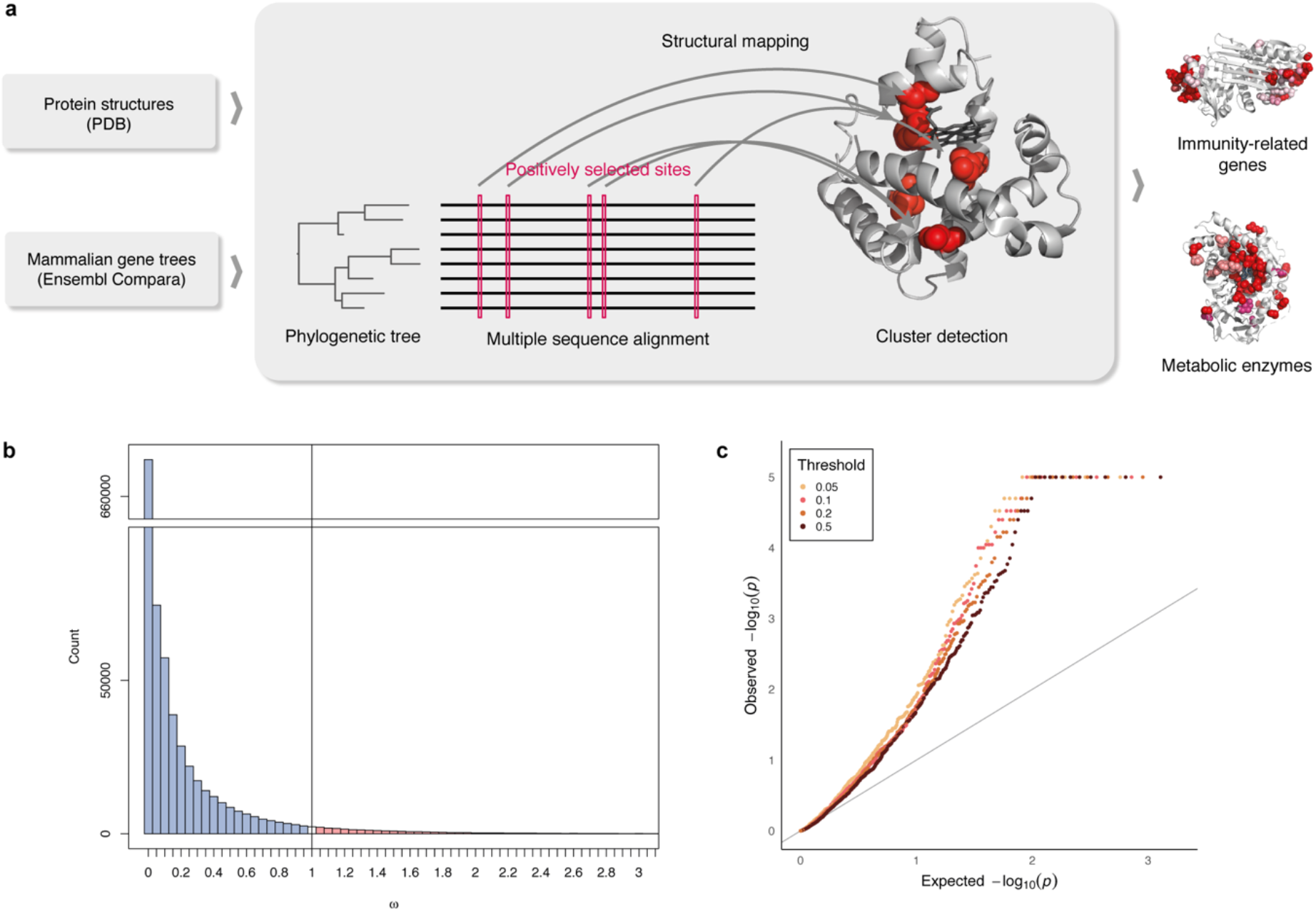
Positively selected residues tend to cluster together. (a) Overview of the approach (b) Distribution of *ω* values in the dataset. With 97.6% of sites having *ω* < 1 (indicating purifying selection), and 2.4% with *ω* ≥ 1, the mean of *ω* across the entire dataset is 0.126. (c) QQ plot of *p*-value distribution obtained from CLUMPS applied to positively selected sites at FDR of 0.05, 0.1, 0.2 and 0.5. If the residues under positive selection were randomly distributed on protein structures, we would expect a uniform distribution of *p*-values (grey line). The observed *p*-values are lower than would be expected under the null hypothesis of random placement, indicating that positively selected sites tend to cluster together.

**Table 1.** Proteins with clusters of positively selected sites. Protein length refers to human orthologs.

| Gene symbol | Gene name | Protein length | PDB | PDB sites | Substrate/relevant ligand in PDB | Number of pos. sel. sites<sup>1</sup> |
| --- | --- | --- | --- | --- | --- | --- |
| **Immune-related proteins** |  |  |  |  |  |  |
| HLA-DRBI | major histocompatibility complex, class II, DR beta 1 | 266 | 1aqd | 187 | endogenous peptide | 16/19/21/25 |
| FCN2 | Ficolin 2 | 313 | 2j3f | 217 | N-acetyl-D-galactosamine | 6/9/10/13 |
| SERPINB3 | Serpin B3 | 390 | 4zk0 | 367 | - | 24/26/31/42 |
| TLR4 | toll-like receptor 4 | 839 | 4g8a | 601 | LPS, LP4 | 51/61/80/119 |
| CD1A | CD1a molecule | 327 | 1onq | 271 | sulfatide self-antigen | 40/44/50/64 |
| C5 | Complement component 5 | 1676 | 3cu7 | 1625 | - | 28/37/53/89 |
| C8A | Complement component 8α | 584 | 3ojy | 478 | - | 16/20/27/34 |
| SIGLEC5 | Sialic acid binding Ig-like lectin 5 | 551 | 2zg1 | 208 | sialic acid | 24/32/38/43 |
| TFRC | transferrin receptor 1 | 760 | 3s9l | 638 | - | 17/19/24/40 |
| **Metabolic enzymes** |  |  |  |  |  |  |
| CYP2C9 | Cytochrome P450, family 2, member C9 | 490 | 1r9o | 453 | flurbiprofen | 29/30/35/40 |
| CYP2D6 | Cytochrome P450, family 2, member D6 | 497 | 2f9q | 454 | - | 6/8/10/14 |
| CYP3A4 | Cytochrome P450, family 3, member A4 | 503 | 3tjs | 449 | desthiazolylmethyloxycarbonyl ritonavir | 22/27/33/41 |
| AKR1B10 | Aldo-keto reductase family 1, member B10 | 316 | 1zua | 316 | tolrestat | 13/18/19/23 |
| AKR1C4 | Aldo-keto reductase family 1, member C4 | 323 | 2fvl | 323 | - | 19/21/23/26 |
| SULT2A1 | Sulfotransferase family 2A member 1 | 285 | 3f3y | 282 | lithocholic acid | 15/18/22/32 |
| CES1 | Carboxylesterase 1 | 568 | 1mx1 | 532 | tacrine | 12/16/19/31 |
| GSTA3 | Glutathione S-transferase alpha 3 | 222 | 1tdi | 218 | glutathione | 11/13/16/20 |
| OLAH | Oleoyl-ACP hydrolase | 318 | 4xjv | 216 | - | 8/12/18/24 |
| AK5 | Adenylate kinase 5 | 562 | 2bwj | 195 | AMP | 3/3/3/3 |
| NCSTN | Nicastrin | 709 | 5a63 | 665 | phosphocholine | 7/8/11/18 |

<sup>1</sup>The number of positively selected sites is given at FDR thresholds of 0.05, 0.1, 0.2 and 0.5, respectively.

## Results

### Identification of positive selection

In order to identify residues that were under positive selection in mammalian evolution, we obtained coding sequences for Eutherian mammals from Ensembl and phylogenetic trees from the Ensembl Compara database ^24^. 3D structures corresponding to human proteins in our dataset were then downloaded from PDB ^25^ and mapped against protein sequences using the SIFTS resource ^26^. We aligned coding sequences corresponding to each tree using the PRANK aligner ^27^ and used the Slr software ^28^ to detect positively selected sites. The resulting dataset comprises 3,347 protein alignments and covers 1,021,133 structure-mapped amino-acid sites. While the majority of sites evolve under purifying selection (fig. 1b), consistent with both theoretical expectations and previous empirical estimates ^13^, we identified 4,498 sites with strong evidence of positive selection (FDR=0.05). We have made these results available as an online resource which allows for displaying and downloading of the structure-mapped sitewise estimates of selective constraint, as well as the underlying alignments and phylogenetic trees (http://www.dev.ebi.ac.uk/goldman-srv/slr/).

### Detecting clustering of positively selected sites

To determine the degree of clustering of positively selected sites, we applied a modification of the CLUMPS algorithm ^21^ to our integrated dataset (see Methods). As the power to detect clustering is limited if very few residues are considered, it is desirable to include as many sites with evidence of positive selection as possible. At the same time, reducing the stringency in the detection of selection by allowing a higher false discovery rate (FDR) can dilute the signal of clustering by including more false positives. As it is not clear *a priori* what the tradeoff between these phenomena is and at what threshold the power to detect clustering is maximised, we applied the chosen clustering detection method separately to positively selected sites detected at different stringency levels. In order to determine the degree to which positively-selected residues form clusters on protein structures, we inspected the overall distribution of *p*-values obtained for each protein from CLUMPS at four FDR thresholds at which positively selected sites were detected (fig. 1c). We find a significant tendency for positively selected sites to cluster together and this trend is maintained at each FDR threshold, indicating that our findings are robust to how stringently positively selected sites are identified.

### Clusters of positively selected sites

Having established that positively selected sites tend to occur close to one another on protein structures, we went on to select cases where evidence for clustering is the strongest. Depending on the FDR threshold used to identify sites as positively selected, between 35 and 52 proteins with clusters of positively selected residues were detected (FDR of clustering <0.05), with substantial overlap between clusters detected at different thresholds (suppl. fig. 2). For 22 proteins, clusters were identified at all four FDR thresholds suggesting that these constitute the most robust findings. For these proteins, we inspected the underlying alignments from which positively selected sites were identified. Correlation on the sequence level can introduce clusters on the level of structure and for this reason it is important to distinguish 3D clusters resulting purely from closeness of sites of interest in the sequence. In all but two cases, we find that positively selected sites are identified in regions of good alignment quality and that clusters of positively selected sites arise mostly from residues that are not adjacent in the sequence and become close to each other only once the protein is folded into its native conformation. The two cases where detected signature of positive selection appears to result from a stretch of contiguous residues in a region of poor alignment quality were rejected from further analysis. The remaining 20 proteins are summarised in table 1. Remarkably, nine of them are immune-related proteins and ten are metabolic enzymes. The remaining protein, nicastrin, is the substrate-recruiting component of gamma-secretase ^29^, a protein complex with catalytic activity and we therefore consider it together with other enzymes.

### Positive selection in proteins involved in immunity

#### Confirmation of validity of clustering approach

Rapid evolutionary rates in genes involved in both adaptive and innate branches of the immune system are a classic example of positive selection ^7,8,30–32^. Proteins where we identified positively-selected clusters (table 1) include cases where positive selection has been documented previously, such as in HLA‑DRB1 ^7^, CD1a ^33^, TLR4 ^33^ and TfR, a protein which is known to have been hijacked by arenaviruses for facilitating cellular entry ^34^. Positively selected residues are located primarily in regions involved in antigen binding, such as the structurally similar binding clefts of HLA-DRB1 and CD1a (suppl. fig. 3–6). While these findings were reported previously, they give confidence in the approach we applied here.

#### Novel findings of selection clusters

We also identify cases where to our knowledge positive selection has not been previously described: ficolin 2 (suppl. fig. 7; though positive selection in the related ficolin-3 has been reported ^37^), complement component 5 (suppl. fig. 8), complement component 8*α* (suppl. fig. 9), SIGLEC5 (suppl. fig. 10) and serpin B3 (fig. 2).

**Figure 2.**
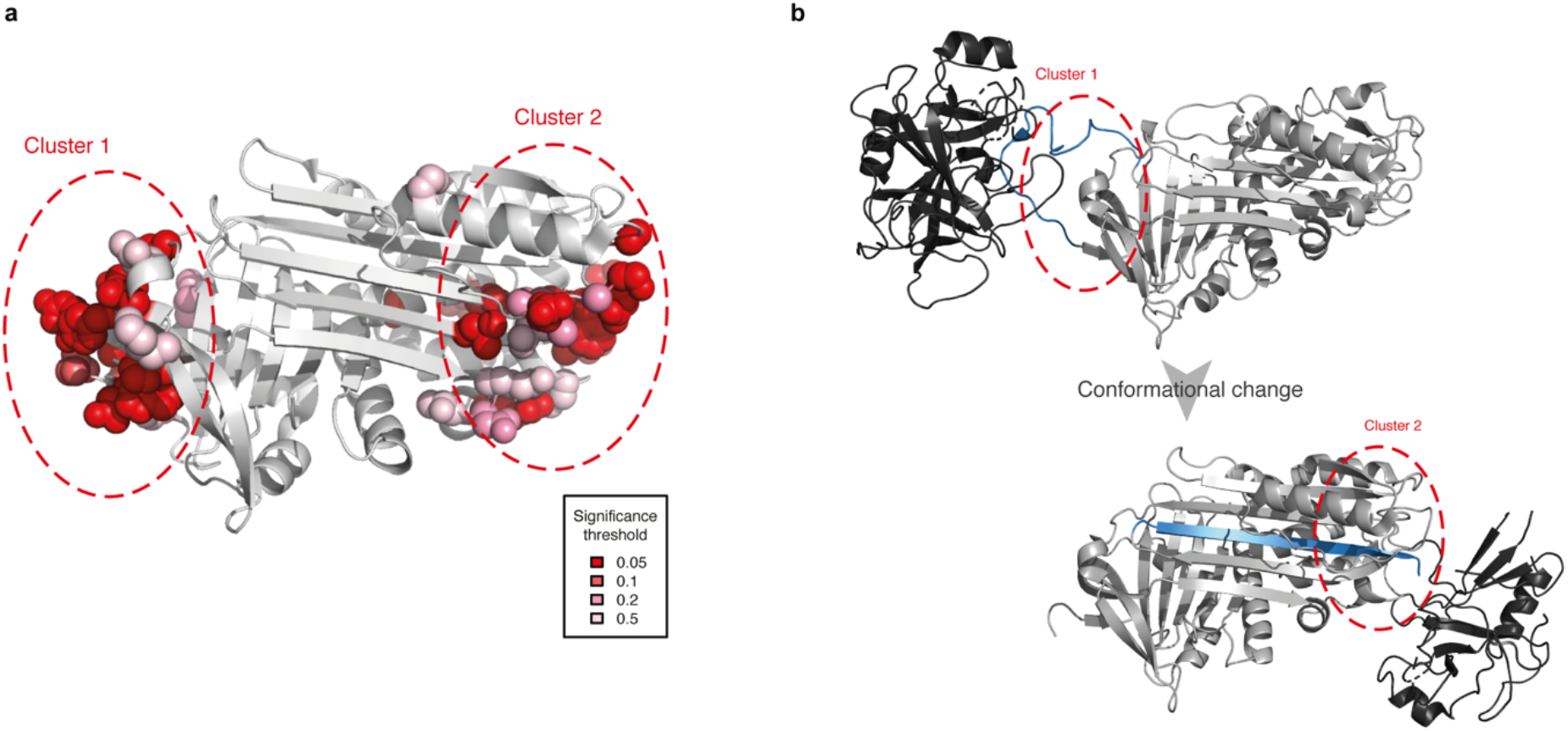
Clusters of positively selected sites in serpin B3. (a) Placement of positively selected sites on the structure of serpin B3 (PDB 4zk0). (b) Mode of action of serpins shown using PDB structures 1k9o (top) and 1ezx (bottom) with the substrate shown in black and the RCL marked in blue. Regions analogous to those where positively selected clusters were marked as in (a). Serpins function by binding their target proteases using a reactive center loop (RCL) that mimics the protease substrate. They then form a covalent bond with the protease and undergo a large conformational change resulting in the protease being deformed and then acylated ^35,36^. We find that positively selected residues surround the RCL and are also located on the opposite side of the protein to which the bound protease is dragged.

The placement of positively selected sites in serpin B3 is particularly interesting as this protein exhibits two clusters concentrated on the opposite poles of the protein (fig. 2a). Serpin B3 belongs to the serpin superfamily of protease inhibitors, though unlike most serpins it binds cysteine rather than serine proteases. Serpins contribute to immunity by inhibiting proteases secreted by bacteria. Serpin B3 inactivates leaked lysosomal cathepsins, inactivates pathogen-derived cathepsins and is also thought to be involved in autoimmunity ^38^. Comparison with other available structures of serpins reveals a remarkable correspondence of these positively selected sites to the protease binding sites before and after the conformational change that characterises the mode of action of serpins (fig. 2b). Furthermore, there is previous evidence that serpin B3 homologs have changed their substrate specificities over the course of evolution consistent with the action of positive selection ^39,40^. The presence of two positively selected residue clusters at opposite poles of the protein implies that both regions participate in the tuning of function. The importance of these regions in the proteolytic function of serpins demonstrates that the positive selection we detect is likely to have functional consequences.

Interactions with pathogens are known to be one of the dominant pressures shaping mammalian evolution ^30^. Our analysis adds mechanistic details to these findings: positively selected clusters in proteins involved in host-pathogen interactions are placed in regions directly mediating binding of pathogen-derived molecules. Binding of pathogen-derived peptides by HLA and subsequent triggering of the immune response is a classic example of this ^41^. Here we have identified further examples of similar mechanisms in components of both innate and adaptive branches of the immune system. Interestingly, these include not only proteins or protein-derived peptides but also lipids (CD1a) and lipopolysaccharides (TLR4). This is true both when binding is facilitating the neutralisation of pathogens and, as in the case of TfR, where host proteins are hijacked by a pathogen to facilitate cellular entry. These scenarios are examples of high evolutionary rate being the result of an ‘arms race’ between host and pathogen. Such dynamics are predicted by the Red Queen hypothesis, which posits that evolution is driven by inter-species competition ^42^.

### Positive selection acting on metabolic enzymes

#### Cytochrome P450s

Ten out of eleven remaining positively selected clusters are found in enzymes. Three of the identified clusters of positively selected sites are in members of the cytochrome P450 (CYP) superfamily (fig. 3). CYPs are the most important drug-metabolising enzyme class, contributing to the metabolism of 90% of drugs as well as many other xenobiotics such as pollutants. These liver enzymes catalyse monooxygenation reactions on a wide range of small and large substrates. More than 50 CYPs have been identified in the human genome but relatively few are known to have a role in drug metabolism ^43^.

**Figure 3.**
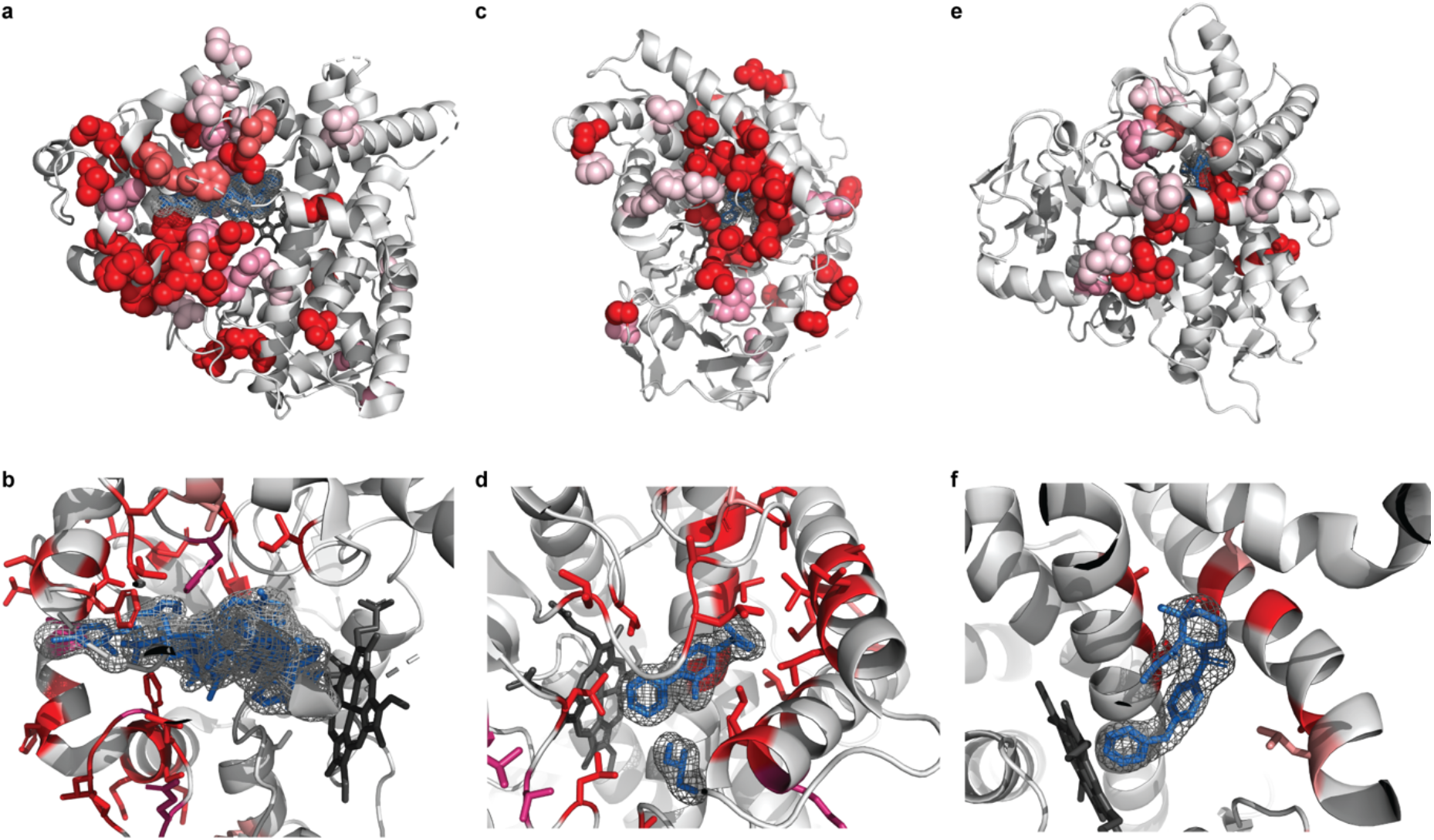
Positively selected residues in CYPs cluster in the substrate entry channel and catalytic site. Positively selected residues in (a, b) CYP3A4 (PDB 3tjs), (c, d) CYP2C9 (PDB 1r9o) and (e, f) CYP2D6 (PDB 2f9q). Hemes are shown coloured in dark grey, other ligands in blue. Additional ligands were transferred from other PDB structures by superimposition: (a, b) desthiazolylmethyloxycarbonyl ritonavir, ketoconazole (PDB 2v0m), erythromycin (PDB 2j0d). (c, d): flurbiprofen (e, f) prinomastat (PDB: 3qm4). Specificity for the extraordinary diversity of substrates in this enzyme superfamily is facilitated by a large, flexible binding pocket at the bottom of which heme is located. In all three structures, the location of the positively selected residues tracks the binding of a ligand, and in general can be found on the sides of helices and in loops that form the binding pocket.

Strikingly, all three of the CYPs where we identify positively selected clusters of residues are known to be important for drug metabolism: CYP3A4 (fig. 3a-b) is the most promiscuous of all CYPs, contributing to the metabolism of ~50% of marketed drugs, and CYP2C9 (fig. 3c-d) and CYP2D6 (fig.3e-f) are also among the six principal CYPs thought to contribute the most to drug metabolism ^44^. In our dataset, alignments containing the three CYPs mentioned before also contain two further cytochrome P450 paralogs that are important for drug metabolism — in total 5 out of 6 enzymes thought to be responsible for the majority of cytochrome P450 drug metabolism show evidence of positive selection.

#### Aldo-keto reductases

We identified positively-selected clusters in two members of the 15 aldo-keto reductases (AKRs) present in human. Similarly to CYPs, AKRs are a family of highly promiscuous enzymes that utilise NAD(P)(H) co-factors and can reduce a wide range of substrates ^45^. AKRs are part of phase II metabolism and can transform or detoxify both endogenous and environmental aldehydes and ketones ^46–48^. Positively selected residues in both AKRs cluster around the region where the substrate binds but not around the NADP+ co-factor (fig.4). This suggests that evolution has tuned substrate specificity while maintaining binding to the cofactor.

**Figure 4.**
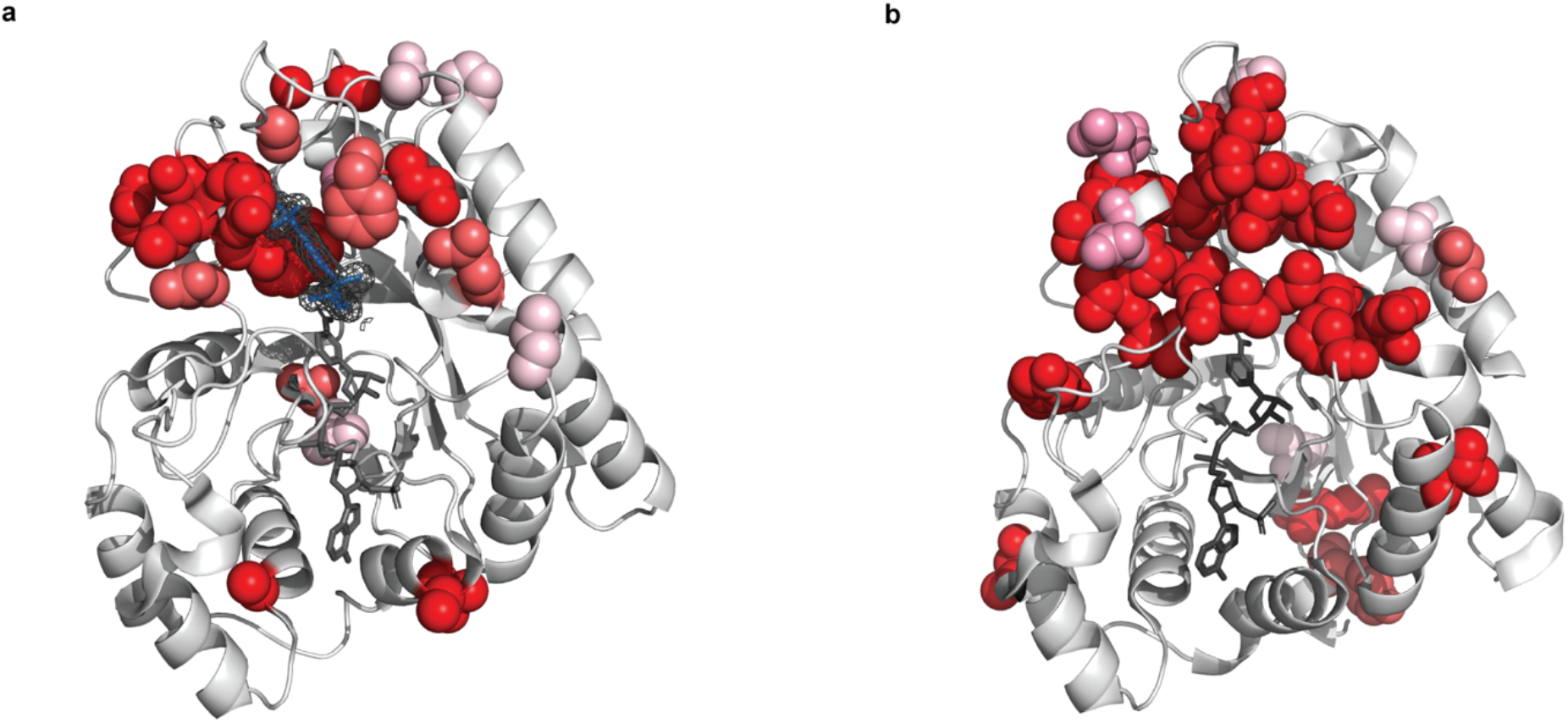
**Positively selected residues in AKRs surround the substrate binding site** positively selected residues in (a) AKR1B10 (PDB 1zua) and (b) AKR1C4 (PDB 2fvl). Tolrestat marked in blue, NADP+ marked in dark grey. Positively selected residues in AKR1B10 cluster around the bound ligand tolrestat, an inhibitor developed for diabetes treatment, but not around the NADP+ cofactor. The structure of AKR1C4 has been solved without ligand but the positively selected residues cluster in a similar region of the structure when compared to AKR1B10. As in the case of AKR1B10, there are no positively selected residues in the neighbourhood of the NADP+ cofactor.

#### Other enzymes

We also identified individual positively selected clusters in the members of three other protein families involved in detoxification: glutathione S-transferase alpha 3 ^49–52^, carboxylesterase 1 ^53,54^ and sulfotransferase 2A1 ^55,56^. In all cases, positively selected sites cluster around the active site of the enzyme where the substrate binds (fig. 5a-c).

**Figure 5.**
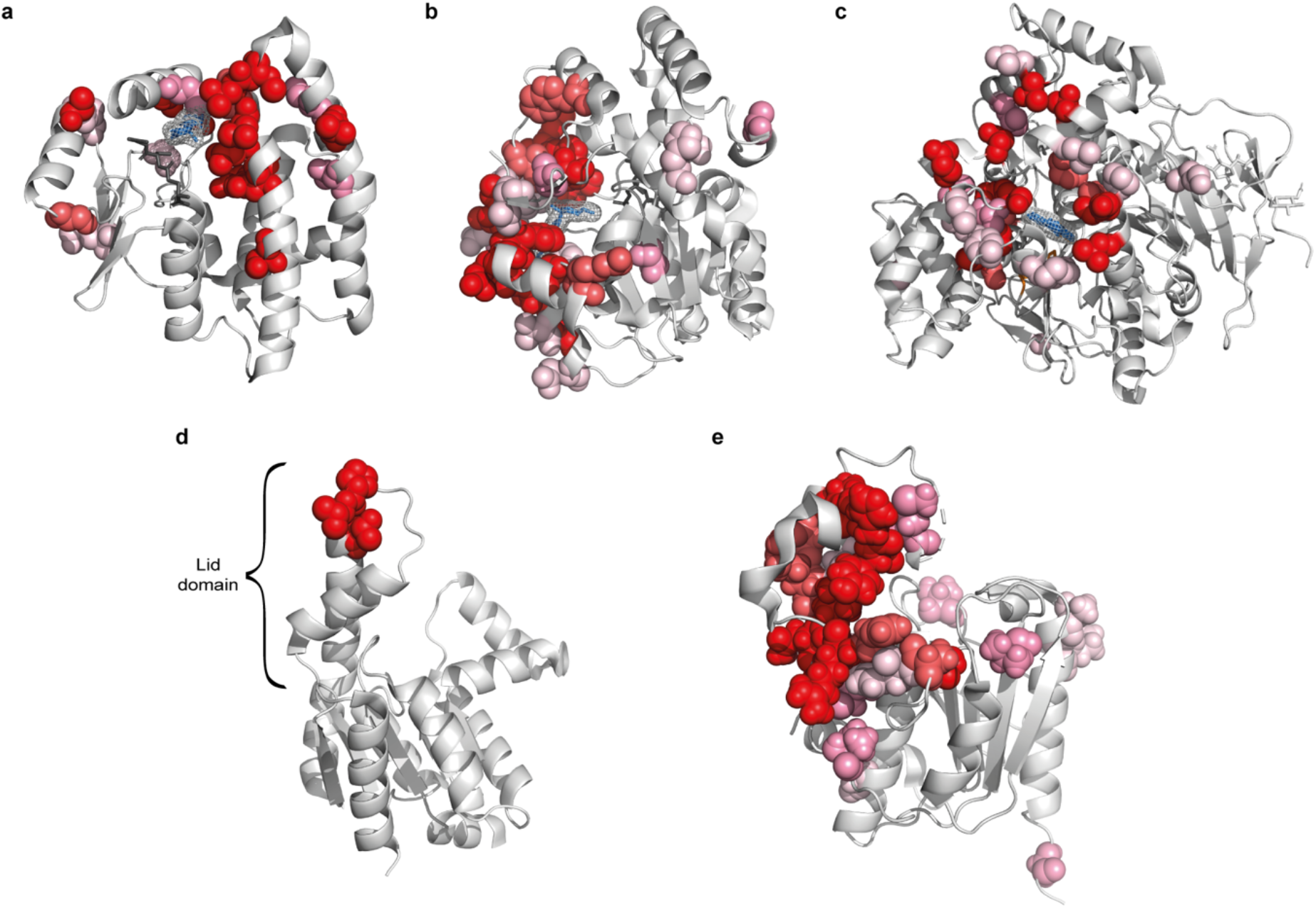
Positively selected residues in other enzymes. (a) Positively selected sites in GSTA3 (PDB 1tdi). Glutathione shown in dark grey, delta-4-androstene-3-17-dione (blue) transferred by structure superimposition from structure 2VCV. (b) Positively selected residues in sulfotransferase 2A1 (PDB 3f3y). Adenosine-3’-5’ diphosphate shown in dark grey, lithocholic acid shown in blue. (c) Positively selected sites in carboxylesterase 1 (PDB 1mx1). Tacrine shown in blue. (d) Positively selected residues in adenylate kinase 5 PDB (PDB 2bwj). (e) Positively selected sites in oleoyl-ACP hydrolase (PDB 4xjv).

In the remaining three cases, positively selected clusters are located in sub-domains that interact with substrates. Adenylate kinase 5 (AK5) is a member of a family of enzymes important for maintaining the energetic balance in the cell by converting ADP into ATP ^57^. Positively selected residues in AK5 fall in the lid sub-domain (fig. 5d) which has been shown to have a role in tuning the enzyme activity ^58^. The three positively selected sites that constitute the positively selected cluster in AK5 flank a DD motif which is highly conserved in AK5 and in other enzymes of the family. Experimentally mutating a residue homologous to V507, one of the sites we have predicted, has been shown to have an effect on the enzyme’s kinetic parameters ^59^, strongly suggesting that the positively selected sites we detect contribute to enzyme specificity and kinetics.

In the case of oleoyl-ACP hydrolase (OLAH), an enzyme involved in controlling the distribution of chain lengths of fatty acids, positively selected residues are located in the capping domain that covers the substrate (fig. 5e). Detailed mutational data for OLAH is lacking but enzymes of the same class have been shown to undergo changes of specificity in other species ^60^. Positively selected sites in nicastrin (suppl. fig. 11) are primarily located in the lid domain that covers the substrate ^61^, and changes at positively selected sites in these enzymes are therefore also consistent with positive selection acting to fine-tune enzymatic activity.

Pervasive positive selection in metabolic enzymes, similar to that experienced by immune-related genes, may seem surprising. Although examples of episodic adaptation of enzymes in specific lineages, particularly in primates, exist ^62,63^, signatures of pervasive positive selection were not previously thought to be common in enzymes. However, enzymes where we identified positively selected clusters are involved in interactions with the environment and share a number of other characteristics that make them plausible targets of positive selection. Eight out of ten such enzymes we identified are involved in the catalysis of xenobiotics. Much like parts of the immune system that directly interact with pathogens, these metabolic enzymes form an interface with the environment and act as one line of defence. The diversification of mammals involved adaptation to varied environments and new diets and as the environment in which they live and feed has changed, so did their exposure to toxins. This is likely to have required widespread, repeated adaptive changes that we observe.

#### Placement of positively-selected sites in relation to functional sites

Having observed the tendency of observed clusters to occur in the direct neighbourhood of bound ligands, we sought to quantify this trend. For structures solved with exogenous ligands, we obtained the distribution of distances for positively-selected residues and compared them to remaining residues (fig. 6a). We find that positively selected residues are significantly closer to those ligands (mean distance 16.9Å vs. 24.4Å; *P* < 2.2×10^−16^; Kolmogorov-Smirnov test), confirming that positively-selected clusters tend to occur closer to bound ligands than would be expected by chance and providing further evidence for positive selection acting to fine-tune ligand binding.

**Figure 6.**
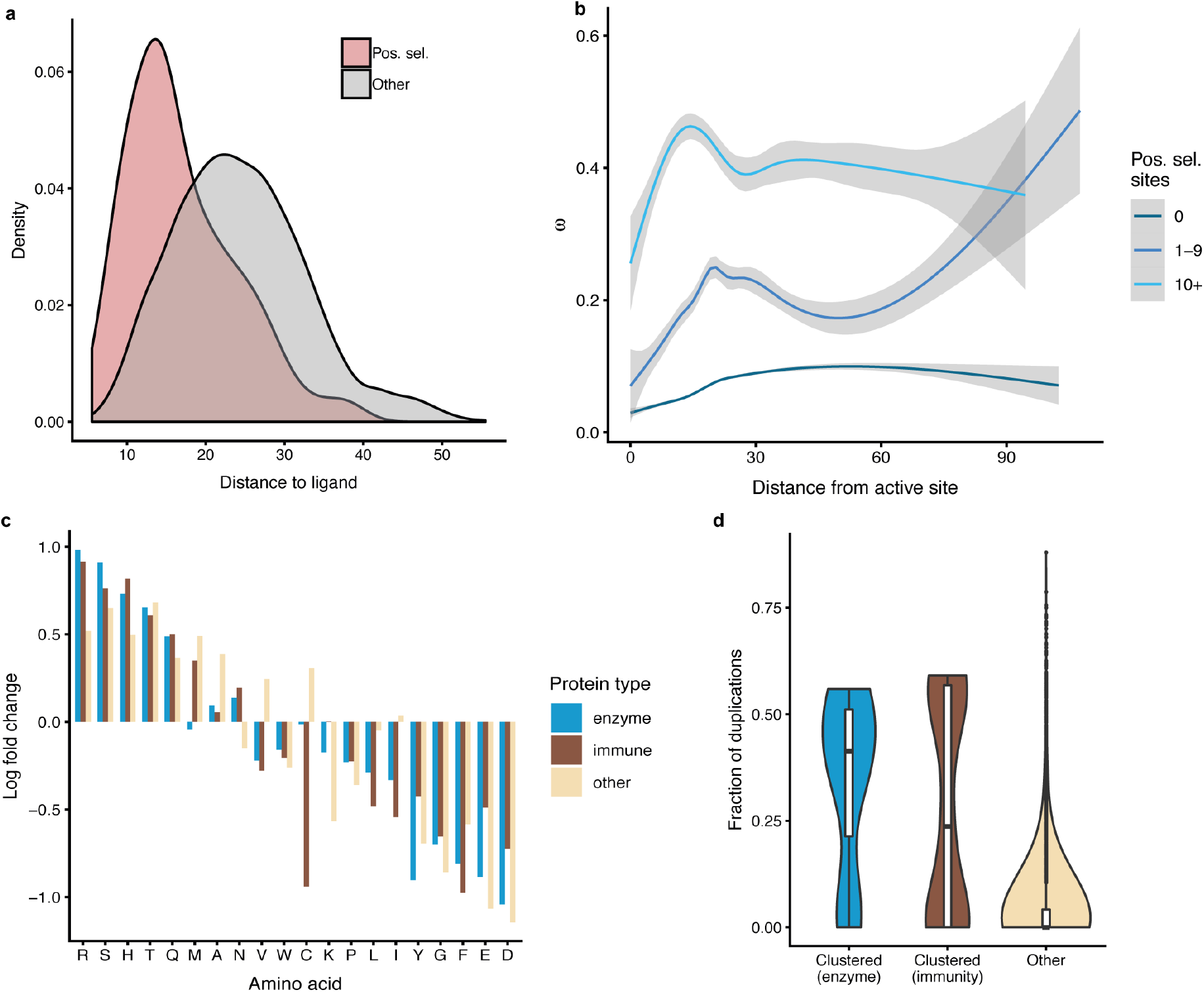
Properties of positively selected sites. (a) Distance of positively selected residues from bound exogenous ligands. (b) The distribution of *ω* as a function of distance from catalytic residues. (c) Departures from the background amino acid frequencies in positively selected residues. (d) Distribution of fraction of gene duplications in proteins with positively selected clusters.

We then investigated the overall distribution of *ω* as a function of distance to catalytic sites, using annotations from Catalytic Site Atlas ^64^. In proteins where we detected no evidence of positive selection, purifying selection is the strongest in the neighbourhood of catalytic sites and gradually relaxes with distance from them (fig. 6b). This trend is consistent with previous studies of selective constraint where positive selection was not considered ^65^. However, in cases where we detected positively selected sites we observed a very different distribution of *ω*, with a peak at 20Å from the catalytic residues. In cases where we detected 10 or more positively selected sites, this trend is even more pronounced, with the peak of *ω* occurring at 14Å from catalytic residues. The enrichment of positively selected residues and elevated mean *ω* in the neighbourhood of catalytic sites indicates that the action of positive selection reshapes the selective constraint on the entire protein structure.

#### Properties of amino acids at positively selected sites

As different regions of proteins are known to have different amino-acid frequencies ^66,67^, we asked whether the positively selected residues we detected exhibit a distinct amino acid distribution. For each protein class, we calculated the change in amino acid frequency at positively selected sites compared to the background frequencies (fig. 6c). While the overall distributions of amino acids are very similar in the different proteins classes (suppl. fig. 12), we observe differences in the distribution of amino acids at positively selected sites compared to the background distribution (fig. 6c). We correlated these enrichment scores with common amino acid physicochemical properties (size, hydrophobicity, net charge and polarity) but found no significant correlations (suppl. table 1), indicating that, while certain amino acids are preferred or avoided at positively selected sites, these trends bear no straightforward relationship to amino acid properties.

#### The role of gene duplication events in adaptive evolution

Gene duplications are thought to be one of the main forces driving evolution, providing ‘raw material’ for evolutionary innovations ^68^. While gene duplications events in themselves are frequently assumed to have no effect on fitness, their retention can be evidence of adaptation ^69^. In order to quantify the effect of duplication events in our dataset, we calculated the fraction of gene duplications (i.e. the number of duplication nodes divided by the total number of nodes) for each phylogenetic tree. We find that both in enzymes and in immune-related genes, the mean paralog fraction is significantly larger than in other genes (0.342 and 0.276, respectively, compared to 0.0397 in the remaining trees; fig. 6d). This trend is significant both in the case of immune proteins and metabolic enzymes (*P* = 0.015 and *P* = 3.1×10^−5^, respectively; Kolmogorov-Smirnov test). This elevated duplication rate in genes where we detected positively selected clusters is consistent with positive selection acting not only on point mutations but also driving gene duplication events to fixation. At the same time, some genes where we detected strong evidence of adaptation (complement component 5, transferrin receptor 1, complement component 8*α*, adenylate kinase 5 and nicastrin) have not undergone any gene duplications, proving that rapid sitewise evolutionary rate and gene duplications can occur independently.

## Discussion

In this study, we curated a dataset covering over one million structurally mapped sites in 3,347 mammalian proteins and assessed the placement of positively selected residues on their 3D structures in an unbiased, genome-wide manner. We find that positively selected sites tend to occur closer to each other in protein structures than is expected by chance and to form clusters in the neighbourhood of functionally important regions. Strikingly, proteins where we found the strongest evidence for clustering of positively selected sites are primarily involved in two major types of environmental responses: host-pathogen interactions and metabolism of xenobiotic compounds. The fact that we observe the strongest evidence of positive selection in these types of proteins gives a coherent view of mammalian evolution being shaped by these two major influences from the environment. Clusters of positively selected sites we identified share both functional and structural similarities and allow us to infer more general principles underlying adaptive evolution.

Xenobiotic-metabolising enzymes are typically able to process a wide range of substrates. Indeed, CYPs and AKRs, where we identified three and two positively selected clusters, respectively, are among the most promiscuous known protein superfamilies. Promiscuous enzymes are thought to be malleable in evolution, as they can maintain their original function as well as acquire specificity for new substrates by going through a promiscuous intermediate which can bind multiple substrates ^70^. The mechanisms by which enzymes acquire new substrates has to date been primarily studied by directed evolution ^71–74^. The examples we have highlighted here provide direct evidence that similar scenarios are also common in natural evolution.

Enzymes involved in xenobiotic metabolism are of great medical relevance, as in humans they are responsible for metabolism of prescribed drugs. Traditional analyses of protein conservation are frequently not suitable for the analysis of genes involved in xenobiotic metabolism, as these tend to evolve rapidly and the analyses used do not explicitly distinguish between neutral evolution and positive selection ^75^. Specific examples we have identified here could be investigated further, for example by detailed mutational studies that have been shown to augment statistical modelling of adaptive evolution ^76^.

Our study highlights the power of incorporating independent sources of information to understand principles governing evolution. The clusters we detected consist of residues that are distributed along the linear sequence of proteins and could not be found without considering protein structure. Consideration of structural information has also allowed us to better understand the mechanistic details of processes underlying adaptation in terms of specific structural and functional features. Information about structural placement of residues can also help to address technical issues that have hindered methods for detecting positive selection. Criticisms levelled at methods for detecting positive selection have revolved around the non-neutrality of synonymous substitutions, local variation in synonymous substitution rate ^77–80^ and the influence of errors in alignment ^81,82^. These phenomena may cause false positives in parts of a protein sequence, but none will result in clustering on protein structure. Structural information can thus serve as an independent validation and a means of demonstrating that observed patterns of positive selection are not a product of confounding factors. Structural clusters can additionally be inspected *post hoc* for proximity to functional features to assess their plausibility and aid interpretation.

The structural and functional similarities we identified here point towards common rules governing the occurrence of pervasive positive selection. Positively selected metabolic enzymes we described here share many structural and functional similarities: positively selected clusters lie in close proximity to bound ligands, indicating that the primary mode in which these enzymes adapt is by affecting residues in the direct neighbourhood of active sites. This finding may seem to contradict the common assumption that functionally important residues are conserved in evolution: for example, the finding that average evolutionary rate is lowest in the neighbourhood of catalytic sites ^65^. However, this is only a superficial disagreement: while functional regions evolve more slowly on average, this does not mean that cannot harbour rapidly evolving, positively selected sites. Indeed, non-functional regions cannot, by definition, undergo adaptive evolution.

As we demonstrate here, while functional regions of proteins are typically more conserved, they can also exhibit a high evolutionary rate that is a hallmark of adaptive evolution. This strongly suggests that instances where positive selection is operating can contradict overall trends of protein evolution. For this reason, it may be counterproductive to incorporate known correlates of evolutionary rate into statistical models for detecting positive selection. In contrast, the fact that positively selected residues can form clusters on protein structures could inform the development of better methods for detecting positive selection. One of the ultimate goals of evolutionary research is integrating evolution of sequence with structure in a general model of protein evolution ^83,84^. Such a universal model of protein evolution has been elusive so far, primarily because the most general approaches require an intractable number of parameters. We would suggest that one way forward is to identify further universal evolutionary trends and gradually incorporate them into mathematical models of protein evolution.

We have demonstrated that analysing selective constraint in the context of structure can help interpret findings and increase their robustness, but all approaches reliant on detailed structural information are limited by the availability and coverage of crystal structures. We hope that the results highlighted here and others we made available online in our web server will assist experimental validation and further understanding of protein function and adaptation. We aimed to establish the relationship between protein structure and the occurrence of positive selection and this proof-of-principle study called for the highest-possible quality data, but incorporating homology-based structural models would be a direct extension to our approach. It is likely that the PDB database is currently biased towards certain protein families which suggests that in future more examples of adaptation will be identified in protein families where currently little or no structural information is available. Similarly, the analysis performed here focused on mammals but could be extended to other clades.

## Acknowledgements

We would like to thank Dr Leo C. James, Dr M. Madan Babu, Dr Patrycja Kozik, Dr Maria Marti-Solano and members of the Goldman Group at EMBL-EBI for helpful discussions and comments on the manuscript.

## Author Contributions

G.S and N.G. conceived the study. G.S. performed all analyses. N.G. supervised the research. G.S. and N.G. wrote the manuscript.

## Competing interests

The authors declare no competing interests.

## Methods

### Genomic data

Coding sequences for mammalian genomes were downloaded from Ensembl ^85^, version 78. Non-eutherian genomes (platypus, gray short-tailed opossum, wallaby and Tasmanian devil) were excluded. Coding sequences for principal isoforms were used. Incomplete and stop codons at ends of sequences were removed.

### Phylogenetic data

The Compara database ^86^ provides gene trees for species stored in Ensembl. The Compara pipeline generates trees containing up to 750 related genes which frequently results in multiple paralogs being included in the same tree. Bearing in mind that selective constraint can be estimated more accurately if more sequences are included, but that including more paralogs can result in averaging over genes which may be under different constraints, we designed a tree-splitting scheme to enable single-gene analysis. As we aimed to maximise the number of orthologous sequences included in each alignment while minimising the number of paralogous sequences, we quantified these criteria in different possible subtrees by calculating the percentage of all species included (taxonomic coverage) and the total number of additional genes for each species beyond the first gene per species (permitting calculation of the paralog fraction). We required a taxonomic coverage of at least 60% and wished to minimise the paralog fraction. To achieve this, starting from each human protein, the tree is traversed towards the root until the desired taxonomic coverage was achieved. Then, the tree is traversed further but only if this does not increase the paralog fraction. The final node of this traversal process and all its descendant nodes then become a tree used for further analysis.

### Sequence alignment

Compara gene trees are reconstructed using principal isoforms and the same sequences were used for alignment. The PRANK aligner ^27^ has been show to limit the number of false positive identifications of positive selection compared to other commonly used aligners ^82,87,88^. PRANK was run in codon mode on sets of sequences corresponding to each Compara-derived tree and with these trees used as guide trees.

### Detecting positive selection

SLR ^28^ was used to obtain sitewise estimates of *ω* within each alignment, using tree topologies from Ensembl Compara and allowing branch lengths to be optimised by SLR. SLR implements a statistical test for positive selection based on the rates of fixation of nonsynonymous and synonymous mutations (*ω*, or dN/dS). This measure is derived from neutral theory ^89,90^, which provides a general framework for studying selective effects and allows for explicitly identifying genomic regions that evolve under positive selection. If mutations that arise are deleterious, they will undergo purifying selection and will be purged from a population, resulting in a low observed evolutionary rate. Conversely, if mutations result in beneficial changes, they will be rapidly driven to fixation. The ratio of fixation probabilities of nonsynonymous and synonymous substitutions can thus be used to estimate the selective constraint acting on the protein level: *ω* ≈ 1 indicates neutral evolution; *ω* < 1 purifying selection; and *ω* > 1 positive selection ^91^. SLR implements the Goldman-Yang codon site model ^92^ similar to that in PAML ^3^. The main difference between SLR and PAML is that SLR makes no assumption about the distribution of *ω* values over the sites of the alignment. SLR first estimates parameters of the phylogenetic model for the entire alignment and then performs a likelihood ratio test between the optimal *ω* and *ω* = 1 for each site. *P*-values reported by SLR associated with each structure-mapped site (see below) were then corrected for multiple testing using the Benjamini-Hochberg FDR method ^93^.

### Structural data

PDB structures matching human proteins in the sequence dataset were downloaded from PDBe ^25^. Structures covering fewer than 100 residues were excluded, and in cases where more than one structure was available the one with the highest sequence similarity to the protein sequence was chosen. In rare cases where more than one human protein with a structure was present for an alignment, one was retained at random. Individual residues were then mapped using the SIFTS database ^26^. SIFTS provides a mapping between PDB ^25^ and UniProt ^94^ sequences and, as the UniProt protein sequences can vary from those in Ensembl, we performed an additional mapping step by constructing pairwise alignments between UniProt and Ensembl sequences, resulting in a sitewise mapping between Ensembl and PDB residues. The pairwise alignments were calculated using the Biopython ^95^ implementation of the Smith-Waterman algorithm ^96^, using the scoring of 1 for matching characters and 0 otherwise, and gap opening and extension penalties of −10 and −0.5 respectively.

### Clustering of positively selected sites

The degree of clustering of the positively selected sites within each protein structure was assessed using the CLUMPS algorithm ^21^. In CLUMPS, the degree of clustering for a set of residues of interest is quantified by the sum of pairwise distances in 3D space. In contrast to the original implementation, we used equal weights for all sites when calculating the pairwise distances. For each set of residues, we then performed 100,000 Monte Carlo simulations permuting the placement of sites by randomly selecting positions from the PDB chain, in order to determine statistical significance of observed patterns. *P*-values resulting from this analysis were then corrected for multiple comparisons using the Benjamini-Hochberg FDR method ^93^.

Statistical analyses were performed in the R environment ^97^.

## References

1. Havrilla, J.M., Pedersen, B.S., Layer, R.M. & Quinlan, A.R. A map of constrained coding regions in the human genome. Nat Genet 51, 88–95 (2019).

2. Fuller, Z.L., Berg, J.J., Mostafavi, H., Sella, G. & Przeworski, M. Measuring intolerance to mutation in human genetics. Nat Genet 51, 772–776 (2019).

3. Yang, Z. PAML 4: phylogenetic analysis by maximum likelihood. Mol Biol Evol 24, 1586–91 (2007).

4. Weaver, S. et al. Datamonkey 2.0: a modern web application for characterizing selective and other evolutionary processes. Mol Biol Evol (2018).

5. Benner, S.A. Natural progression. Nature 409, 459 (2001).

6. Echave, J., Spielman, S.J. & Wilke, C.O. Causes of evolutionary rate variation among protein sites. Nat. Rev. Genet. 17, 109–121 (2016).

7. Hughes, A.L. & Nei, M. Pattern of nucleotide substitution at major histocompatibility complex class I loci reveals overdominant selection. Nature 335, 167–70 (1988).

8. Sawyer, S.L., Wu, L.I., Emerman, M. & Malik, H.S. Positive selection of primate TRIM5alpha identifies a critical species-specific retroviral restriction domain. Proc Natl Acad Sci U S A 102, 2832–7 (2005).

9. Schott, R.K., Refvik, S.P., Hauser, F.E., Lopez-Fernandez, H. & Chang, B.S. Divergent positive selection in rhodopsin from lake and riverine cichlid fishes. Mol Biol Evol 31, 1149–65 (2014).

10. Endo, T., Ikeo, K. & Gojobori, T. Large-scale search for genes on which positive selection may operate. Mol Biol Evol 13, 685–90 (1996).

11. Kosiol, C. et al. Patterns of positive selection in six Mammalian genomes. PLoS Genet 4, e1000144 (2008).

12. Eory, L., Halligan, D.L. & Keightley, P.D. Distributions of selectively constrained sites and deleterious mutation rates in the hominid and murid genomes. Mol Biol Evol 27, 177–92 (2010).

13. Lindblad-Toh, K. et al. A high-resolution map of human evolutionary constraint using 29 mammals. Nature 478, 476–82 (2011).

14. Roux, J. et al. Patterns of positive selection in seven ant genomes. Mol Biol Evol 31, 1661–85 (2014).

15. Cicconardi, F., Marcatili, P., Arthofer, W., Schlick-Steiner, B.C. & Steiner, F.M. Positive diversifying selection is a pervasive adaptive force throughout the Drosophila radiation. Mol Phylogenet Evol 112, 230–243 (2017).

16. Yang, Z., Nielsen, R. & Goldman, N. In defense of statistical methods for detecting positive selection. Proc Natl Acad Sci U S A 106, E95; author reply E96 (2009).

17. Zhai, W., Nielsen, R., Goldman, N. & Yang, Z. Looking for Darwin in genomic sequences--validity and success of statistical methods. Mol Biol Evol 29, 2889–93 (2012).

18. Laskowski, R.A. & Thornton, J.M. Understanding the molecular machinery of genetics through 3D structures. Nat Rev Genet 9, 141–51 (2008).

19. Homburger, J.R. et al. Multidimensional structure-function relationships in human beta-cardiac myosin from population-scale genetic variation. Proc Natl Acad Sci U S A 113, 6701–6 (2016).

20. Miller, M.L. et al. Pan-Cancer Analysis of Mutation Hotspots in Protein Domains. Cell Syst 1, 197–209 (2015).

21. Kamburov, A. et al. Comprehensive assessment of cancer missense mutation clustering in protein structures. Proc Natl Acad Sci U S A 112, E5486–95 (2015).

22. Niu, B. et al. Protein-structure-guided discovery of functional mutations across 19 cancer types. Nat Genet 48, 827–37 (2016).

23. Araya, C.L. et al. Identification of significantly mutated regions across cancer types highlights a rich landscape of functional molecular alterations. Nat Genet 48, 117–25 (2016).

24. Vilella, A.J. et al. EnsemblCompara GeneTrees: Complete, duplication-aware phylogenetic trees in vertebrates. Genome Res 19, 327–35 (2009).

25. Mir, S. et al. PDBe: towards reusable data delivery infrastructure at protein data bank in Europe. Nucleic Acids Res 46, D486–D492 (2018).

26. Dana, J.M. et al. SIFTS: updated Structure Integration with Function, Taxonomy and Sequences resource allows 40-fold increase in coverage of structure-based annotations for proteins. Nucleic Acids Res 47, D482–D489 (2019).

27. Löytynoja, A. & Goldman, N. Phylogeny-aware gap placement prevents errors in sequence alignment and evolutionary analysis. Science 320, 1632–1635 (2008).

28. Massingham, T. & Goldman, N. Detecting amino acid sites under positive selection and purifying selection. Genetics 169, 1753–62 (2005).

29. Xie, T. et al. Crystal structure of the gamma-secretase component nicastrin. Proc Natl Acad Sci U S A 111, 13349–54 (2014).

30. Enard, D., Cai, L., Gwennap, C. & Elife, D.A.P. Viruses are a dominant driver of protein adaptation in mammals. cdn.elifesciences.org.

31. Webb, A.E. et al. Adaptive Evolution as a Predictor of Species-Specific Innate Immune Response. Mol Biol Evol 32, 1717–29 (2015).

32. Ebel, E.R., Telis, N., Venkataram, S., Petrov, D.A. & Enard, D. High rate of adaptation of mammalian proteins that interact with Plasmodium and related parasites. PLOS Genetics 13, e1007023 (2017).

33. Sironi, M., Cagliani, R., Forni, D. & Clerici, M. Evolutionary insights into host-pathogen interactions from mammalian sequence data. Nat Rev Genet 16, 224–36 (2015).

34. Demogines, A., Abraham, J., Choe, H., Farzan, M. & Sawyer, S.L. Dual host-virus arms races shape an essential housekeeping protein. PLoS Biol. 11, e1001571 (2013).

35. Huntington, J.A., Read, R.J. & Carrell, R.W. Structure of a serpin-protease complex shows inhibition by deformation. Nature 407, 923–926 (2000).

36. Ye, S. et al. The structure of a Michaelis serpin-protease complex. Nat. Struct. Biol. 8, 979–983 (2001).

37. Kim, P.M., Korbel, J.O. & Gerstein, M.B. Positive selection at the protein network periphery: evaluation in terms of structural constraints and cellular context. Proc. Natl. Acad. Sci. U. S. A. 104, 20274–20279 (2007).

38. Vidalino, L. et al. SERPINB3, apoptosis and autoimmunity. Autoimmun Rev 9, 108–12 (2009).

39. Heit, C. et al. Update of the human and mouse SERPIN gene superfamily. Hum. Genomics 7, 22 (2013).

40. Izuhara, K., Ohta, S., Kanaji, S., Shiraishi, H. & Arima, K. Recent progress in understanding the diversity of the human ov-serpin/clade B serpin family. Cell Mol Life Sci 65, 2541–53 (2008).

41. Hughes, A.L., Ota, T. & Nei, M. Positive Darwinian selection promotes charge profile diversity in the antigen-binding cleft of class I major-histocompatibility-complex molecules. Mol Biol Evol 7, 515–24 (1990).

42. Van Valen, L. A new evolutionary law, 1—30 (1973).

43. Lynch, T. & Price, A. The effect of cytochrome P450 metabolism on drug response, interactions, and adverse effects. Am Fam Physician 76, 391–6 (2007).

44. Wilkinson, G.R. Drug metabolism and variability among patients in drug response. N Engl J Med 352, 2211–21 (2005).

45. Penning, T.M. The aldo-keto reductases (AKRs): Overview. Chem Biol Interact 234, 236–46 (2015).

46. Jin, Y. & Penning, T.M. Aldo-keto reductases and bioactivation/detoxication. Annu Rev Pharmacol Toxicol 47, 263–92 (2007).

47. Bachur, N.R. Cytoplasmic aldo-keto reductases: a class of drug metabolizing enzymes. Science 193, 595–7 (1976).

48. Barski, O.A., Tipparaju, S.M. & Bhatnagar, A. The aldo-keto reductase superfamily and its role in drug metabolism and detoxification. Drug Metab. Rev. 40, 553–624 (2008).

49. Gloss, A.D. et al. Evolution in an ancient detoxification pathway is coupled with a transition to herbivory in the drosophilidae. Mol. Biol. Evol 31, 2441–2456 (2014).

50. Lan, T., Wang, X.-R. & Zeng, Q.-Y. Structural and functional evolution of positively selected sites in pine glutathione S-transferase enzyme family. J. Biol. Chem. 288, 24441–24451 (2013).

51. da Fonseca, R.R., Johnson, W.E., O’Brien, S.J., Vasconcelos, V. & Antunes, A. Molecular evolution and the role of oxidative stress in the expansion and functional diversification of cytosolic glutathione transferases. BMC Evol. Biol. 10, 281 (2010).

52. Ivarsson, Y., Mackey, A.J., Edalat, M., Pearson, W.R. & Mannervik, B. Identification of Residues in Glutathione Transferase Capable of Driving Functional Diversification in Evolution: A novel approach to protein redesign. J. Biol. Chem. 278, 8733–8738 (2003).

53. Wang, D. et al. Human carboxylesterases: a comprehensive review. Acta Pharm Sin B 8, 699–712 (2018).

54. Bencharit, S., Morton, C.L., Xue, Y., Potter, P.M. & Redinbo, M.R. Structural basis of heroin and cocaine metabolism by a promiscuous human drug-processing enzyme. Nat Struct Biol 10, 349–56 (2003).

55. Allali-Hassani, A. et al. Structural and chemical profiling of the human cytosolic sulfotransferases. PLoS Biol 5, e97 (2007).

56. Gamage, N. et al. Human sulfotransferases and their role in chemical metabolism. Toxicol Sci 90, 5–22 (2006).

57. Kerns, S.J. et al. The energy landscape of adenylate kinase during catalysis. Nat. Struct. Mol. Biol. 22, 124–131 (2015).

58. Schrank, T.P., Wrabl, J.O. & Hilser, V.J. Conformational Heterogeneity Within the LID Domain Mediates Substrate Binding to Escherichia coli Adenylate Kinase: Function Follows Fluctuations. in Topics in Current Chemistry (eds. Klinman, J. & Schiffer, S.H.) 95–121 (Springer Berlin Heidelberg, 2013).

59. Schrank, T.P., Wrabl, J.O. & Hilser, V.J. Conformational heterogeneity within the LID domain mediates substrate binding to Escherichia coli adenylate kinase: function follows fluctuations. Top Curr Chem 337, 95–121 (2013).

60. Jing, F. et al. Phylogenetic and experimental characterization of an acyl-ACP thioesterase family reveals significant diversity in enzymatic specificity and activity. BMC Biochem. 12, 44 (2011).

61. Bai, X.C. et al. An atomic structure of human gamma-secretase. Nature 525, 212–217 (2015).

62. Messier, W. & Stewart, C.B. Episodic adaptive evolution of primate lysozymes. Nature 385, 151–4 (1997).

63. Zhang, J., Zhang, Y.P. & Rosenberg, H.F. Adaptive evolution of a duplicated pancreatic ribonuclease gene in a leaf-eating monkey. Nat Genet 30, 411–5 (2002).

64. Furnham, N. et al. The Catalytic Site Atlas 2.0: cataloging catalytic sites and residues identified in enzymes. Nucleic Acids Res 42, D485–9 (2014).

65. Jack, B.R., Meyer, A.G., Echave, J. & Wilke, C.O. Functional Sites Induce Long-Range Evolutionary Constraints in Enzymes. PLoS Biol 14, e1002452 (2016).

66. Goldman, N., Thorne, J.L. & Jones, D.T. Assessing the impact of secondary structure and solvent accessibility on protein evolution. Genetics 149, 445–58 (1998).

67. Bartlett, G.J., Porter, C.T., Borkakoti, N. & Thornton, J.M. Analysis of Catalytic Residues in Enzyme Active Sites. J. Mol. Biol. 324, 105–121 (2002).

68. Ohno, S. Evolution by gene duplication, xv, 160 p. (Springer-Verlag, London, 1970).

69. Francino, M.P. An adaptive radiation model for the origin of new gene functions. Nat Genet 37, 573–7 (2005).

70. Khersonsky, O., Roodveldt, C. & Tawfik, D.S. Enzyme promiscuity: evolutionary and mechanistic aspects. Curr Opin Chem Biol 10, 498–508 (2006).

71. Schmidt, D.M. et al. Evolutionary potential of (beta/alpha)8-barrels: functional promiscuity produced by single substitutions in the enolase superfamily. Biochemistry 42, 8387–93 (2003).

72. Rothman, S.C. & Kirsch, J.F. How does an enzyme evolved in vitro compare to naturally occurring homologs possessing the targeted function? Tyrosine aminotransferase from aspartate aminotransferase. J Mol Biol 327, 593–608 (2003).

73. Hoffmeister, D., Yang, J., Liu, L. & Thorson, J.S. Creation of the first anomeric D/L-sugar kinase by means of directed evolution. Proc Natl Acad Sci U S A 100, 13184–9 (2003).

74. Aharoni, A. et al. The ‘evolvability’ of promiscuous protein functions. Nat Genet 37, 73–6 (2005).

75. Zhou, Y., Mkrtchian, S., Kumondai, M., Hiratsuka, M. & Lauschke, V.M. An optimized prediction framework to assess the functional impact of pharmacogenetic variants. Pharmacogenomics J 19, 115–126 (2019).

76. Bloom, J.D. Identification of positive selection in genes is greatly improved by using experimentally informed site-specific models. Biol. Direct 12, 1 (2017).

77. Parmley, J.L., Chamary, J.V. & Hurst, L.D. Evidence for purifying selection against synonymous mutations in mammalian exonic splicing enhancers. Mol Biol Evol 23, 301–9 (2006).

78. Macossay-Castillo, M., Kosol, S., Tompa, P. & Pancsa, R. Synonymous constraint elements show a tendency to encode intrinsically disordered protein segments. PLoS Comput Biol 10, e1003607 (2014).

79. Savisaar, R. & Hurst, L.D. Both Maintenance and Avoidance of RNA-Binding Protein Interactions Constrain Coding Sequence Evolution. Mol Biol Evol 34, 1110–1126 (2017).

80. Davydov, II, Salamin, N. & Robinson-Rechavi, M. Large-Scale Comparative Analysis of Codon Models Accounting for Protein and Nucleotide Selection. Mol Biol Evol (2019).

81. Schneider, A. et al. Estimates of positive Darwinian selection are inflated by errors in sequencing, annotation, and alignment. Genome Biol Evol 1, 114–8 (2009).

82. Jordan, G. & Goldman, N. The effects of alignment error and alignment filtering on the sitewise detection of positive selection. Mol Biol Evol 29, 1125–39 (2012).

83. Perron, U., Moal, I., Thorne, J. & Goldman, N. Probabilistic Models for the Study of Protein Evolution. In D. Balding, I. Moltke, and J. Marioni, editors, Handbook of Statistical Genetics. Wiley-Interscience, 4th edition. (In Press).

84. Perron, U., Kozlov, A.M., Stamatakis, A., Goldman, N. & Moal, I.H. Modelling structural constraints on protein evolution via side-chain conformational states. Mol. Biol. Evol. (In Press).

## References

85. Cunningham, F. et al. Ensembl 2019. Nucleic Acids Res 47, D745–D751 (2019).

86. Herrero, J. et al. Ensembl comparative genomics resources. Database 2016(2016).

87. Fletcher, W. & Yang, Z. The effect of insertions, deletions, and alignment errors on the branch-site test of positive selection. Mol Biol Evol 27, 2257–67 (2010).

88. Markova-Raina, P. & Petrov, D. High sensitivity to aligner and high rate of false positives in the estimates of positive selection in the 12 Drosophila genomes. Genome Res 21, 863–74 (2011).

89. Kimura, M. Evolutionary rate at the molecular level. Nature 217, 624–6 (1968).

90. Kimura, M. On the probability of fixation of mutant genes in a population. Genetics 47, 713–9 (1962).

91. Nei, M. & Gojobori, T. Simple methods for estimating the numbers of synonymous and nonsynonymous nucleotide substitutions. Mol Biol Evol 3, 418–26 (1986).

92. Goldman, N. & Yang, Z. A codon-based model of nucleotide substitution for protein-coding DNA sequences. Mol Biol Evol 11, 725–36 (1994).

93. Benjamini, Y. & Hochberg, Y. Controlling the False Discovery Rate: A Practical and Powerful Approach to Multiple Testing. Journal of the Royal Statistical Society. Series B (Methodological) 57, 289–300 (1995).

94. UniProt Consortium, T. UniProt: the universal protein knowledgebase. Nucleic Acids Res 46, 2699 (2018).

95. Cock, P.J. et al. Biopython: freely available Python tools for computational molecular biology and bioinformatics. Bioinformatics 25, 1422–3 (2009).

96. Smith, T.F. & Waterman, M.S. Identification of common molecular subsequences. J Mol Biol 147, 195–7 (1981).

97. Development Core Team, R. R Core Team. R: A Language and Environment for Statistical Computing 2014. (2013).

