## Supplementary Material for "Integrated evolutionary and structural analysis reveals xenobiotics and pathogens as the major drivers of mammalian adaptation"

### Supplementary figures

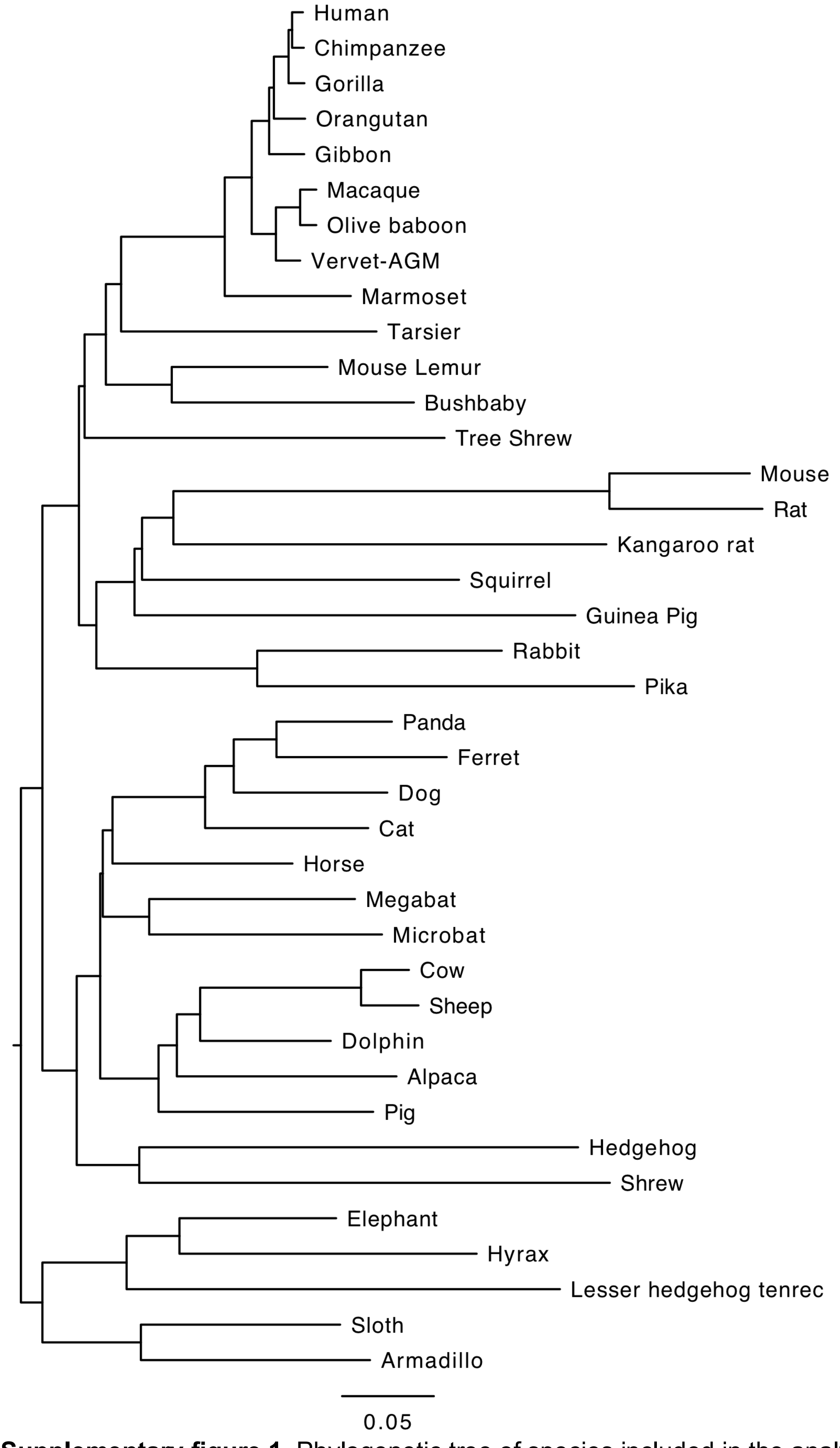


**Supplementary figure 1.** Phylogenetic tree of species included in the analysis.

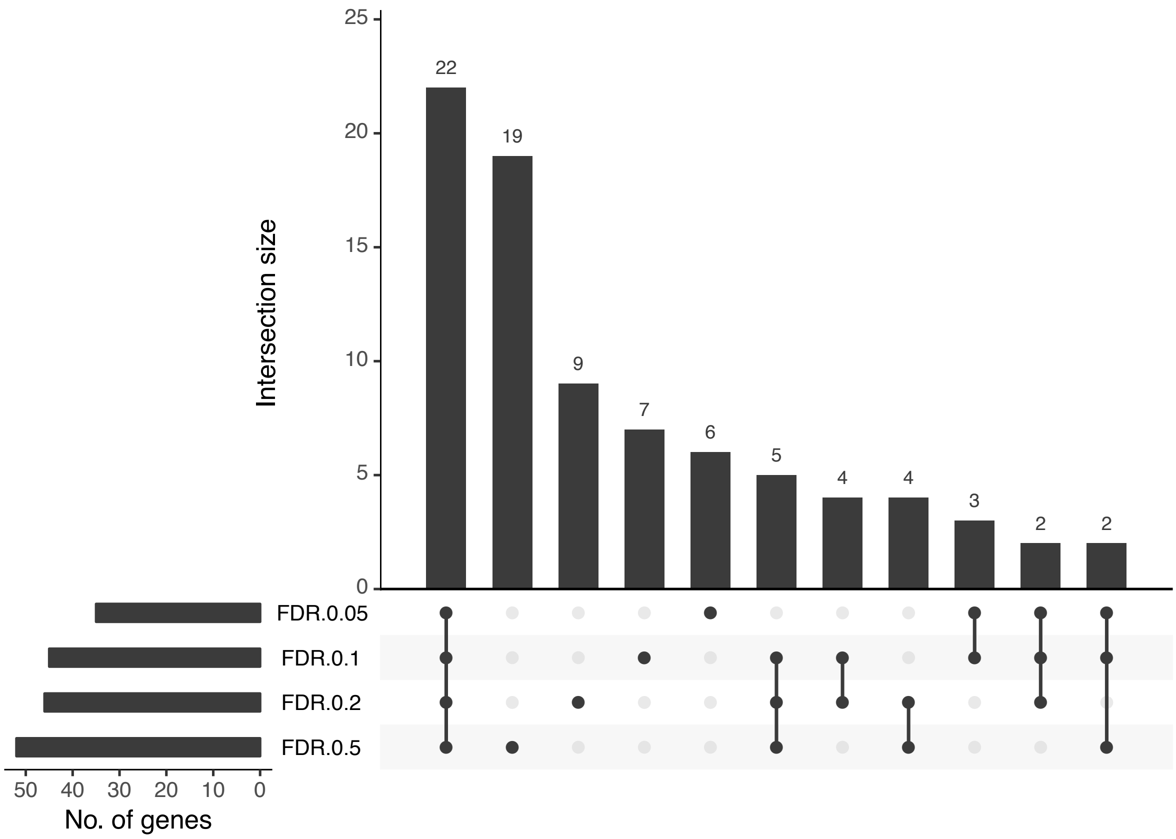


**Supplementary figure 2.** Overlap between detected clusters of positively selected residues between four levels of stringency at which positively selected residues are identified.

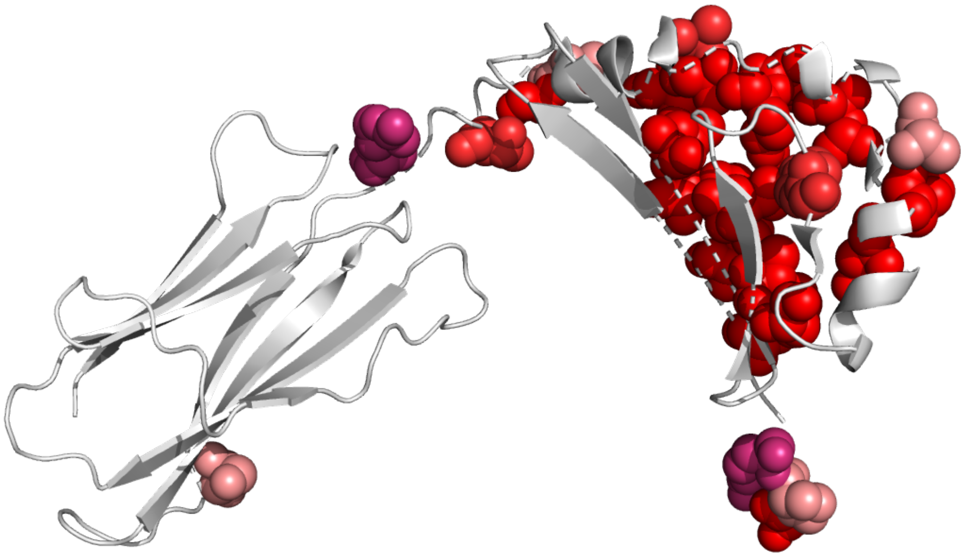

**Supplementary figure 3.** Positively selected sites in major histocompatibility complex class II (PDB: 1aqd).

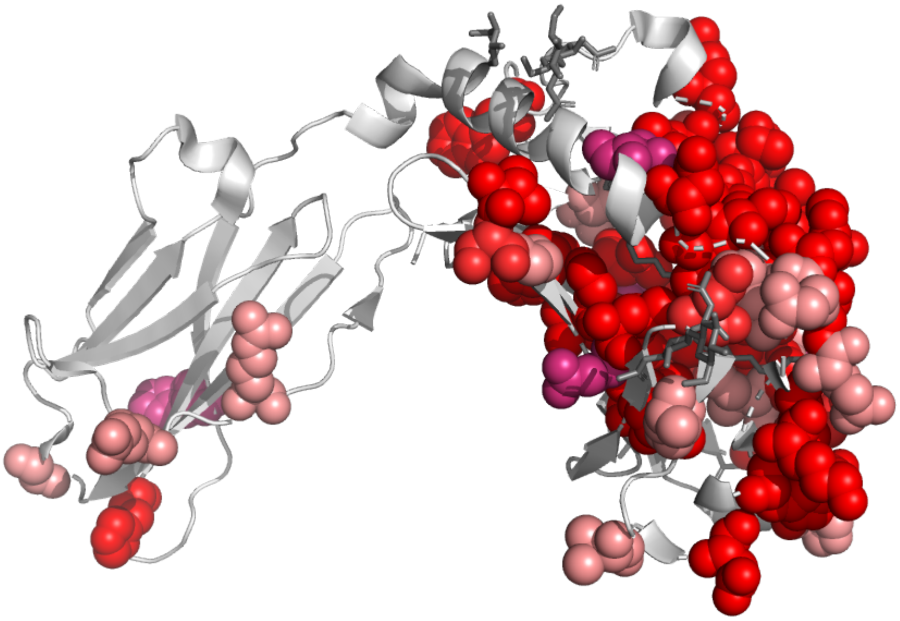

**Supplementary figure 4.** Positively selected sites in CD1a molecule (PDB: 1onq).

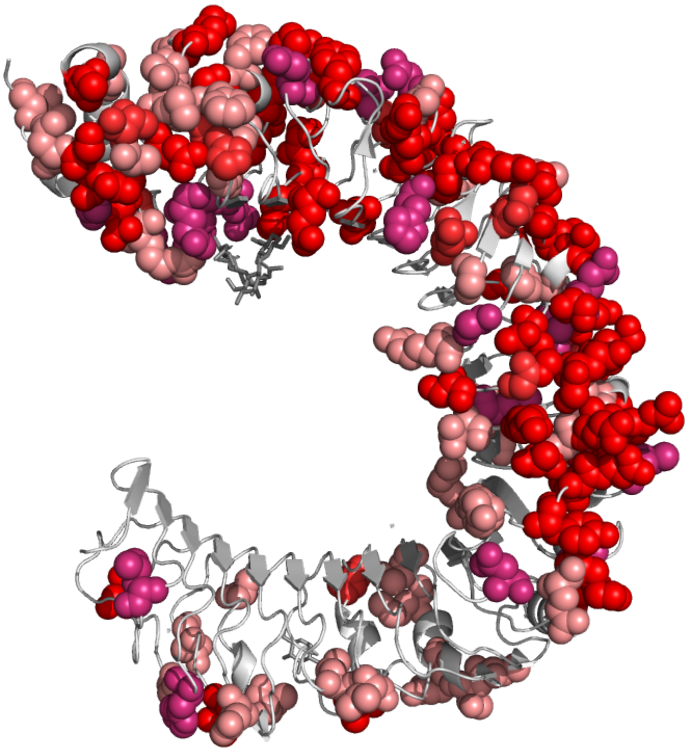

**Supplementary figure 5.** Positively selected sites in toll-like receptor 4 (PDB: 4g8a).

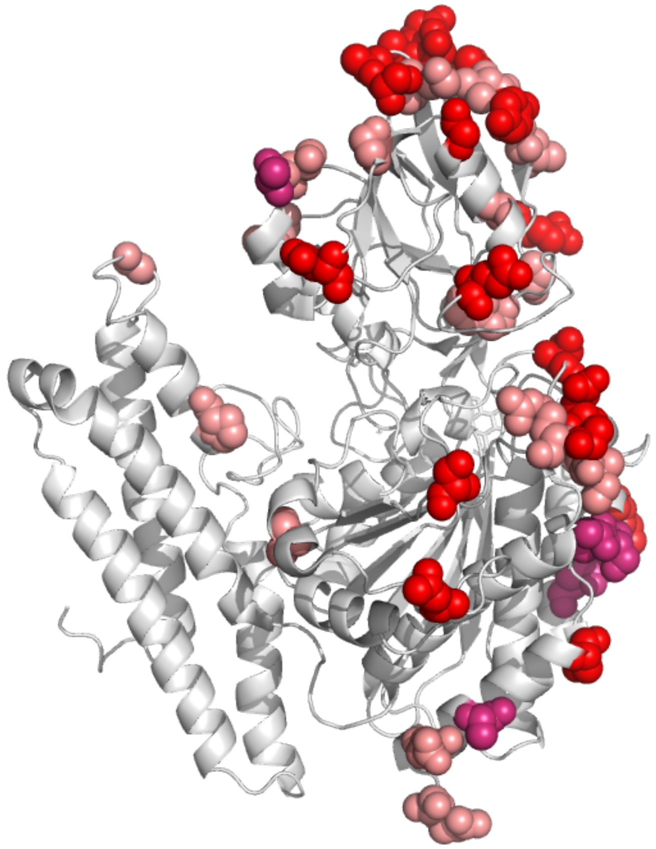

**Supplementary figure 6.** Positively selected sites in transferrin receptor 1 (PDB: 3s9l).

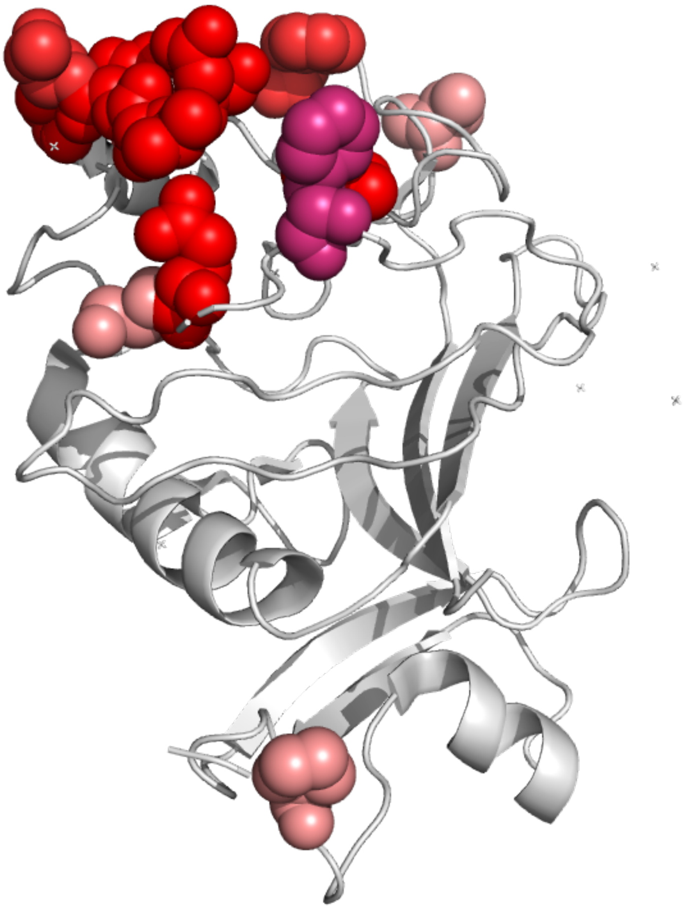

**Supplementary figure 7.** Positively selected sites in ficolin 2 (PDB: 2j3f).

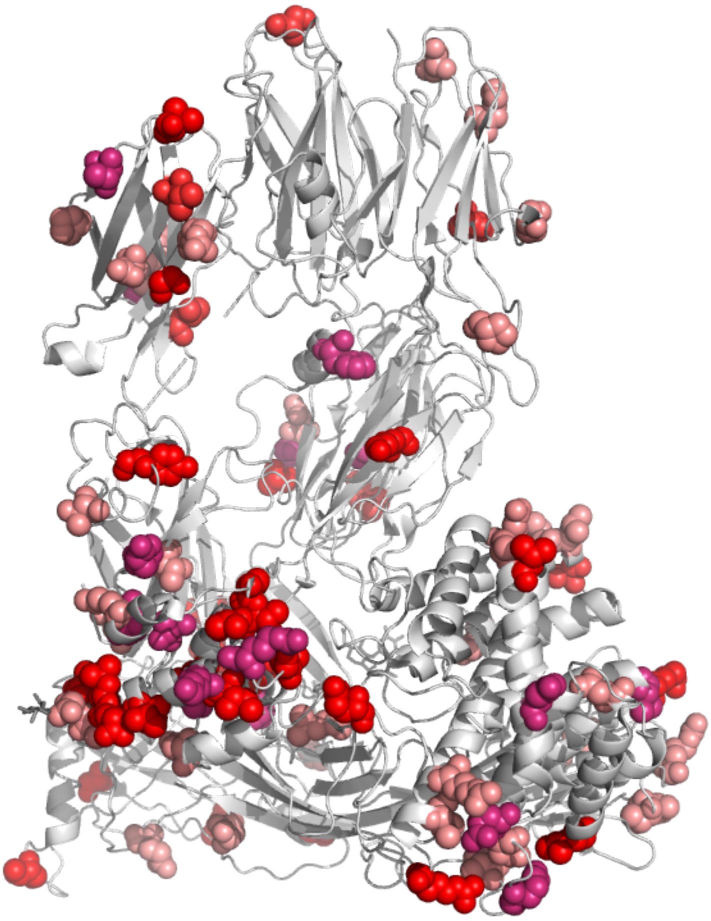

**Supplementary figure 8.** Positively selected sites in complement component C5 (PDB: 3cu7).

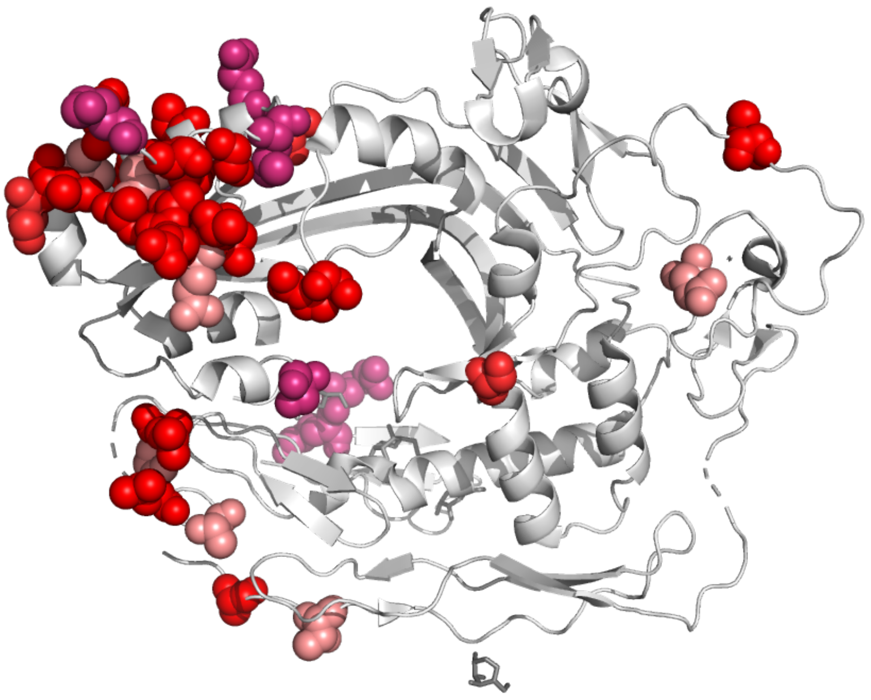

**Supplementary figure 9.** Positively selected sites in complement component C8 (PDB: 3ojy).

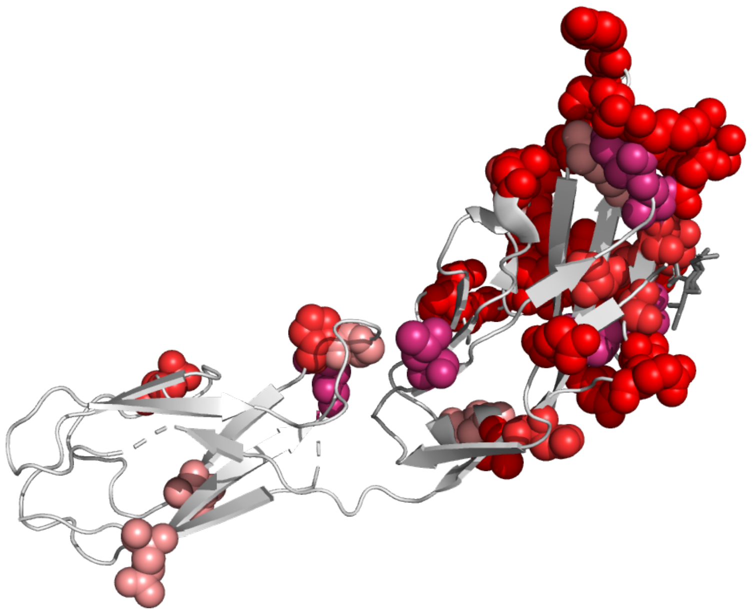

**Supplementary figure 10.** Positively selected sites in siglec-5 (PDB: 2zg1).

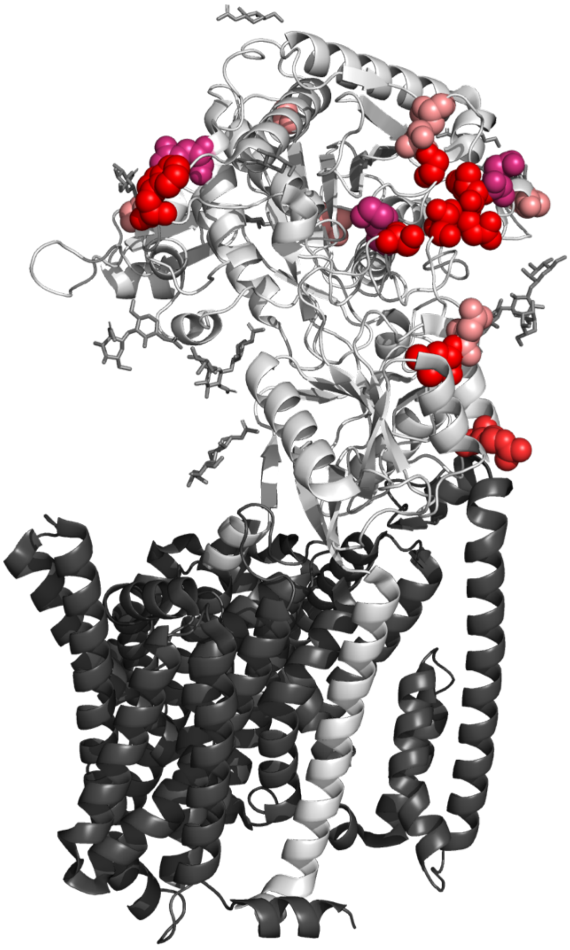

**Supplementary figure 11.** Positively selected sites in nicastrin (PDB: 5a63).

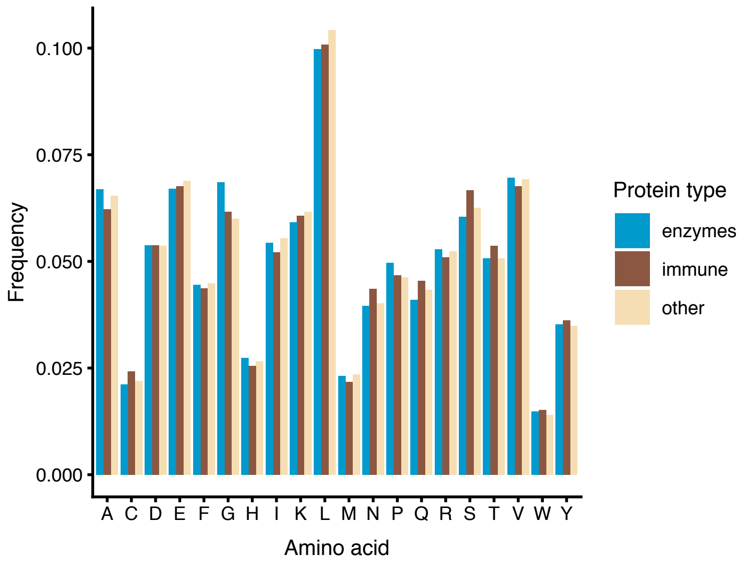


**Supplementary figure 12.** Amino acid frequencies in enzymes, immune-related proteins and remaining proteins.

| Property | Pearson's r | P-value | Adj. p-value |
|---|---|---|---|
| Size (Dawson, 1972) | 0.211 | 0.373 | 0.932 |
| Hydrophobicity (Levitt, 1976) | 0.067 | 0.780 | 0.941 |
| Net charge (Klein et al., 1984) | 0.460 | 0.042 | 0.207 |
| Polarity (Zimmerman et al., 1968) | -0.018 | 0.942 | 0.941 |

**Supplementary table 1.** Correlations between amino-acid physicochemical properties and frequency changes at positively selected residues.
